# Advancing Snow Leopard Density Estimates to Assess Population Dynamics Over Time in Afghanistan

**DOI:** 10.1101/2025.08.25.672058

**Authors:** Eve Bohnett, Sorosh Poya Faryabi, Rebecca Lewison, Li An, Stephane Ostrowski

## Abstract

Effective wildlife monitoring requires innovative methodologies to ensure accurate population assessments for conservation planning. This study focuses on snow leopards (Panthera uncia), a species classified as Vulnerable by the IUCN due to habitat loss, climate change, and over-exploitation. Employing spatial capture-recapture (SCR) models and AI-assisted individual identification, we analyzed camera trapping data from 2012-2013 and 2017-2019 to assess changes in snow leopard density and abundance in Wakhan National Park, Afghanistan. Our analysis used single-session and multi-session models to provide a comprehensive view of population dynamics. Single-session models revealed a substantial increase in density, with the full model estimate rising from 1.38 ± 0.39 SE to 3.57 ± 0.88 SE, while right-only and left-only models showed increases from 1.43 ± 0.53 SE to 3.38 ± 1.01 SE and from 1.04 ± 0.36 SE to 2.21 ± 0.72 SE, respectively. Our findings also highlight the importance of incorporating bilateral asymmetry into population estimates to avoid overestimation. For single-session models, the full model density estimate was 11.7% higher in 2012-2013 and 27.8% higher in 2017-2019 compared to the average of the left-only and right-only estimates. For multi-session models, the full model density estimate was 13.0% higher in 2012-2013 and 46.9% higher in 2017-2019. This emphasizes the importance of addressing asymmetry to avoid potential overestimation. This study demonstrates the potential of combining advanced analytical frameworks with AI to improve wildlife population assessments. By refining SCR methodologies and addressing key biases, these methods provide critical insights for snow leopard conservation and broader applications to wildlife monitoring. While the results indicate a significant increase in snow leopard populations, continued monitoring and long-term studies are essential to account for changes due to environmental and anthropogenic factors.

## Introduction

Estimating population density and size is a crucial element of applied ecology, but it presents considerable difficulties for cryptic terrestrial carnivores, such as snow leopards, that exist at low numbers (Sharma et al. 2020a). Snow leopards are distributed in low densities over large areas of high-altitude habitat (Sharma et al. 2020a; Solari et al. 2023), necessitating large sampling areas to produce statistically robust population estimates (Jackson et al. 2006; Alexander et al. 2016). The global assessment of snow leopards has relied on density estimates from less than 3% of the snow leopard’s range (Sharma and Singh 2021), and reported densities in the literature vary widely from 1.46 - 8.48 animals per 100 km^2^ (Jackson et al. 2006; Alexander et al. 2015, 2016; Zhang et al. 2020). Although these estimates are realistically thought to be within 0.9-1.8 individuals per 100 km^2^ (McCarthy et al. 2016), discrepancies are influenced by various factors. To collect appropriate data to determine the sizes and densities of snow leopard populations, traditional, yet less reliable, pugmark-based classification systems (Riordan 1998) have been mainly replaced by photographic capture-recapture analysis (Jackson et al. 2006; Alexander et al. 2015) or genetic analysis of fecal scats (Janečka et al. 2011; Laguardia et al. 2015). Among these, camera trap photographic sampling has been widely used, allowing for individual mark-recapture events across multiple camera trap stations to estimate the population size or density in a given area (Jackson et al. 2006; Alexander et al. 2015; Zhang et al. 2020; Sharma et al. 2020b). Recent studies indicate that global and local population estimates from camera trapping are likely inflated due to misidentification in camera trap data and sampling biases in areas with higher densities (Ellis 2018; Johansson et al. 2020). This highlights the need for improved data sampling and processing workflows to produce more accurate population estimates.

Camera trapping methods have increasingly required advanced statistical techniques and diverse parameterizations, with spatial capture-recapture (SCR) emerging as the most prominent approach (Efford et al. 2009; Royle et al. 2014; Sutherland et al. 2019). While snow leopard density estimates vary due to biological and environmental factors—such as seasonal movements, breeding behavior, and habitat suitability (Jackson et al. 2006; Janečka et al. 2011) —several data processing and modeling workflow steps can introduce biases that should be mitigated. SCR relies on identifying individuals based on unique traits, such as pelage patterns or morphological features, which can introduce biases that compromise the accuracy of population density estimates. These include (1) classification errors during individual identification (Choo et al. 2020; Johansson et al. 2020; Bohnett et al. 2023b), (2) accounting for misidentification errors due to unpaired camera stations capturing one side of animals with bilateral asymmetry (McClintock et al. 2013; Augustine et al. 2018; Maronde et al. 2020), and (3) sorting data by demographic factors such as sex (Sollmann et al. 2011; Xiao et al. 2016; Augustine et al. 2019). Additionally, SCR model parameters related to sampling characteristics, such as spatial extent and detection distances, can significantly affect density estimates (Keiter et al. 2017).

A major source of error arises from observer misclassification of camera trap images, where the skill of the classifier impacts the precision of individual identification (Johansson et al. 2020; Bohnett et al. 2023b). Recent studies on snow leopard identification have demonstrated that Whiskerbook.org, an artificial intelligence (AI) software program, can assist “expert” manual observers in identifying individuals with minimal classification errors (Bohnett et al. 2023b). Although AI has not yet been fully applied to snow leopard density estimation, its use has shown potential in reducing misclassifications, thereby improving population size and density estimates.

Even with AI-assisted classification, misidentification errors remain possible, particularly due to the natural bilateral asymmetry of snow leopard markings when only one flank is captured by unpaired camera stations. This asymmetry complicates individual identification, as a single snow leopard may be erroneously classified as two individuals if only left-flank images are captured in one instance and right-flank images in another (Augustine et al. 2018). One potential solution is to classify individual encounters by flank (left-only, right-only, or both) and select the side with the most captures for analysis (Wilson et al. 1999; Wang and Macdonald 2009; Berrow 2012; Nair et al. 2012; Srivathsa et al. 2015). Another approach is integrated SCR modeling of photo encounters (left, right, both) modeling (McClintock et al. 2013) or employing a “spatial partial identity model (SPIM)” framework (Augustine et al. 2018). Few snow leopard studies have used paired camera setups to capture both flanks simultaneously (Jackson et al. 2006; Sharma et al. 2020b), likely due to the cost, making unpaired camera setups more common. Some researchers, such as Pal et al. (2022), have restricted analyses to individuals captured on both flanks or those with extensive captures on one side, similar to methods applied to other species (Wang and Macdonald 2009; Kalle et al. 2011; Nair et al. 2012). However, this approach can introduce bias depending on whether left or right-flank data are preferentially analyzed. Positive bias arises when datasets with more individuals or captures are selected, while negative bias occurs when captures disproportionately represent individuals with higher detection probabilities. Most snow leopard density studies have overlooked differences between left- and right-flank data, leading to potential biases and overestimations of population size. However, researchers can reduce the risk of biased estimates by analyzing flank-specific datasets separately, integrating them, or utilizing partial identity models.

The absence of any methodology to address bilateral asymmetry in snow leopard density studies highlights the need for more robust and informed approaches, especially given the species’ low densities. Addressing these challenges requires researchers to adopt rigorous methods that account for observer bias and misidentification. Evaluating approaches is essential for improving snow leopard population estimates across their range.

This study utilized camera trap data collected between 2012-2013 and 2017-2019 to assess changes in snow leopard density in the Hindukush Mountains of Wakhan National Park, Afghanistan. Individual identification was performed using an artificial intelligence software, Whiskerbook, to assist expert manual observers in classifying individual snow leopards based on photographic captures (Bohnett et al. 2023b, a). The analysis employed spatial capture-recapture (SCR) models, focusing on the bilateral asymmetry of snow leopard markings by analyzing left-only and right-only capture histories. The approach addresses a common limitation in snow leopard density studies, where capture histories typically combine left- and right-sided images, potentially introducing bias. By comparing these separate capture histories, this study highlights the impact of bilateral asymmetry on density estimates and the importance of addressing this issue in population assessments.

Furthermore, the study explored the application of single-session and multi-session open spatial capture-recapture (oSCR) models to evaluate their influence on density estimates, reliability, and potential biases. The findings contribute to refining methodological approaches in snow leopard population studies by providing insights into the limitations of existing workflows, including misidentification errors and biases introduced by unpaired camera stations. This work emphasizes the need for rigorous and standardized approaches to improve the accuracy of snow leopard population estimates across their range. The results are discussed in the broader context of snow leopard conservation, highlighting the challenges and opportunities in safeguarding this iconic species in Wakhan National Park and other parts of its range.

### Methods Study Area

Wakhan National Park (36.8454° N, 72.2837° E) covers 10,950 km^2^ in Afghanistan’s Badakhshan Province, bordering Tajikistan, Pakistan, and China (Smallwood and Shank 2019) (Figure 1). The park is approximately 350 km long and 13-65 km wide, situated at the geological intersection of the Hindukush, Pamirs, and Karakoram Mountain ranges. Managed as an IUCN Category VI protected area, it encompasses 70% of Afghanistan’s confirmed snow leopard habitat (Moheb and Paley 2016). Threats, such as poaching and retaliatory killing, have prompted the development of new levels of local governance and protection, with the support of the Wildlife Conservation Society (WCS) since 2006.

**Figure 1:**
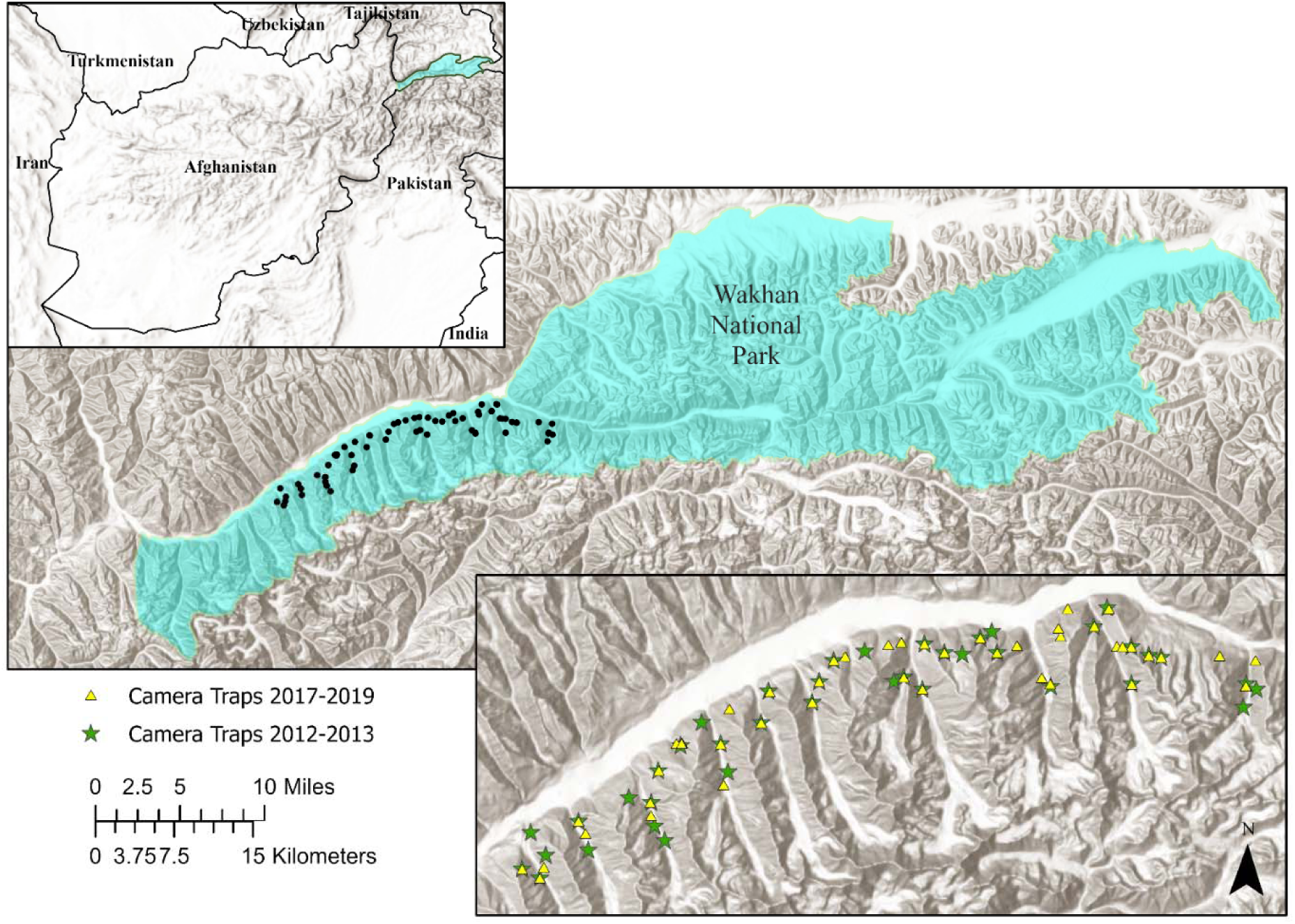
A map of Wakhan National Park, Badakhshan Province, Afghanistan, with the location of camera traps mapped in the Hindu Kush region of the Wakhan National Park.

### Camera-Trap Methods

Camera traps were deployed across ravines and ridges to sample the study area. Between October 2012 and June 2013, WCS Afghanistan installed 43 camera stations (Reconyx and Bushnell), averaging 34 cameras per two-month session. Camera spacing ranged from 1,033 m to 6,039 m, with an average of 2,221 m, and the traps were placed at an average elevation of 3,681 m (min: 2,786 m, max: 4,441 m). In a second deployment between October 2017 and May 2018, 67 camera stations were installed, averaging 33 cameras per session, at elevations of 3,681 m (min: 3,081 m, max: 4,481 m), and distances between cameras ranged from 767 m to 3,894 m (average 2,013 m). A third deployment took place between June and October 2019, with WCS deploying 30 camera traps across the same study area.

Previous GPS collaring studies estimated snow leopard home ranges between 26 km² and 395 km² (Rosenbaum et al. 2023), with a mean of 207 km² ± 63 km² for males and 124 km² ± 41 km² for females (Johansson et al. 2016). Camera trap design recommendations suggest a dispersed setup with camera spacing of at least one per home range, ideally at a magnitude larger than the home range but less than 8 km (Nawaz et al. 2021).

### Individual Identification Methods

We organized snow leopard photographs into “encounters,” defined as successive images of individual snow leopards separated by at least one hour. Each encounter was manually reviewed to identify individuals, and cloud-based batch processing of camera trap data was conducted using Whiskerbook.org, a program developed by Wild Me (Berger-Wolf et al. 2017). Whiskerbook employs two algorithms—pose-invariant embeddings (PIE) and HotSpotter—to provide reliable matching of individual snow leopards (Bohnett et al. 2023a). Studies have demonstrated that the Whiskerbook is less prone to error compared to manual sorting (Bohnett et al. 2023b). In our study, we used the Whiskerbook to classify the imagery using the AI tools Hotspotter and PIE (Bohnett et al. 2023a), along with a manual “expert” observer to further qualify the images using the visual matcher tool in the Whiskerbook program. In our study, the software was used with expert manual classification using Whiskerbook’s visual matcher tool. Extensive rounds of sorting ensured the highest accuracy by verifying three consistent and three inconsistent markings on the snow leopard’s pelage for individual identification (Sharma et al. 2020b; Johansson et al. 2020).

### Quality Control

After the initial identification of individuals, each encounter was manually labeled for quality (Low, High) in a final dataset, which underwent an additional quality control process. Encounters were labeled based on whether the animal was captured facing left, right, or both sides. Low-quality photos, defined by blurriness, poor lighting, unfavorable angles, or severe glare, were excluded from the analysis. Further checks included reviewing distances between capture locations to verify consistency in individual identifications, particularly for encounters recorded at substantial distances apart. All individual IDs were finally rigorously reviewed three times for consistency across encounters. Additionally, 27 juveniles, representing 35 encounters, were removed from the dataset before analysis, as juveniles do not yet establish home ranges. These steps were implemented to increase the certainty of individual identifications and minimize potential bias.

### Spatial Capture-Recapture (SCR) Methods

Spatial capture-recapture (SCR) models estimate animal movement and space use by incorporating camera trap locations and detection probabilities. The design optimizes camera placement based on study area size and individual species’ movement patterns (Sun et al. 2014). SCR models are particularly advantageous for species like the snow leopard, as they do not require sampling grids larger than the area occupied by the species (Sollmann et al. 2012). For this study, we collated data on individual snow leopard encounters at each camera station within 24-hour intervals for analysis. Population density and abundance were estimated using single-session SCR models implemented with the oSCR package in R (Royle et al. 2009; Efford and Fewster 2013; Borchers and Fewster 2016; Efford 2021b; Sutherland 2022). The models also assess the probability of detecting individuals based on activity center distance.

Surveys from 2012-2013 (Block 1) were divided into six 64–65-day sessions, and surveys from 2017-2019 (Block 2) were divided into six 63–68-day sessions (Table 1).

**Table 1.** The survey sessions used in the single and multi-session oSCR analysis include session start / end dates, number of days, and deployed cameras.

| Session | Block | Start | End | Season | Days | Cameras |
| --- | --- | --- | --- | --- | --- | --- |
| 1 | 1 | 2012-10-13 | 2012-12-16 | Winter | 65 | 28 |
| 2 | 1 | 2012-12-17 | 2013-02-19 | Winter | 65 | 32 |
| 3 | 1 | 2013-02-20 | 2013-04-24 | Spring | 64 | 38 |
| 4 | 1 | 2013-04-25 | 2013-06-28 | Spring | 65 | 43 |
| 5 | 1 | 2013-06-29 | 2013-09-01 | Summer | 65 | 39 |
| 6 | 1 | 2013-09-02 | 2013-11-05 | Fall | 65 | 28 |
| 7 | 2 | 2017-09-18 | 2017-11-18 | Fall | 63 | 34 |
| 8 | 2 | 2017-11-19 | 2018-01-20 | Winter | 63 | 34 |
| 9 | 2 | 2018-01-21 | 2018-03-25 | Winter | 64 | 34 |
| 10 | 2 | 2018-03-26 | 2018-05-27 | Spring | 63 | 34 |
| 11 | 2 | 2019-06-21 | 2019-08-27 | Summer | 68 | 28 |
| 12 | 2 | 2019-08-28 | 2019-11-03 | Fall | 68 | 30 |

### Buffer and Buffer Masks

In oSCR models, buffer masks around sampling areas represent the “area of integration” where animals are expected to be detected (Sollmann et al. 2012; Sun et al. 2014). Buffer size was initially set based on half the minimum distance moved (½ MMDM) to estimate sigma, reflecting each session’s average home range size (Sollmann et al. 2012; Sun et al. 2014). The buffer size and resolution were calibrated using the oSCR package in R, with preliminary adjustments to understand how density estimates responded to different buffer sizes (Zimmermann and Foresti 2016). Final buffer sizes were determined by multiplying sigma by a factor of four, following calibration and literature guidance (Royle et al. 2014). The resulting buffer covered approximately 20,000 meters, with an 800-meter spatial grid resolution. Further filters were applied, including an altitude cutoff of 5,600 meters was applied to the buffer, based on telemetry data from 2012-2014, which tracked four snow leopards in the study area. The buffer had a hard boundary in the north due to the Panj and Lower Pamir Rivers, as no evidence suggested snow leopards crossed these rivers (Figure 3).

**Figure 3:**
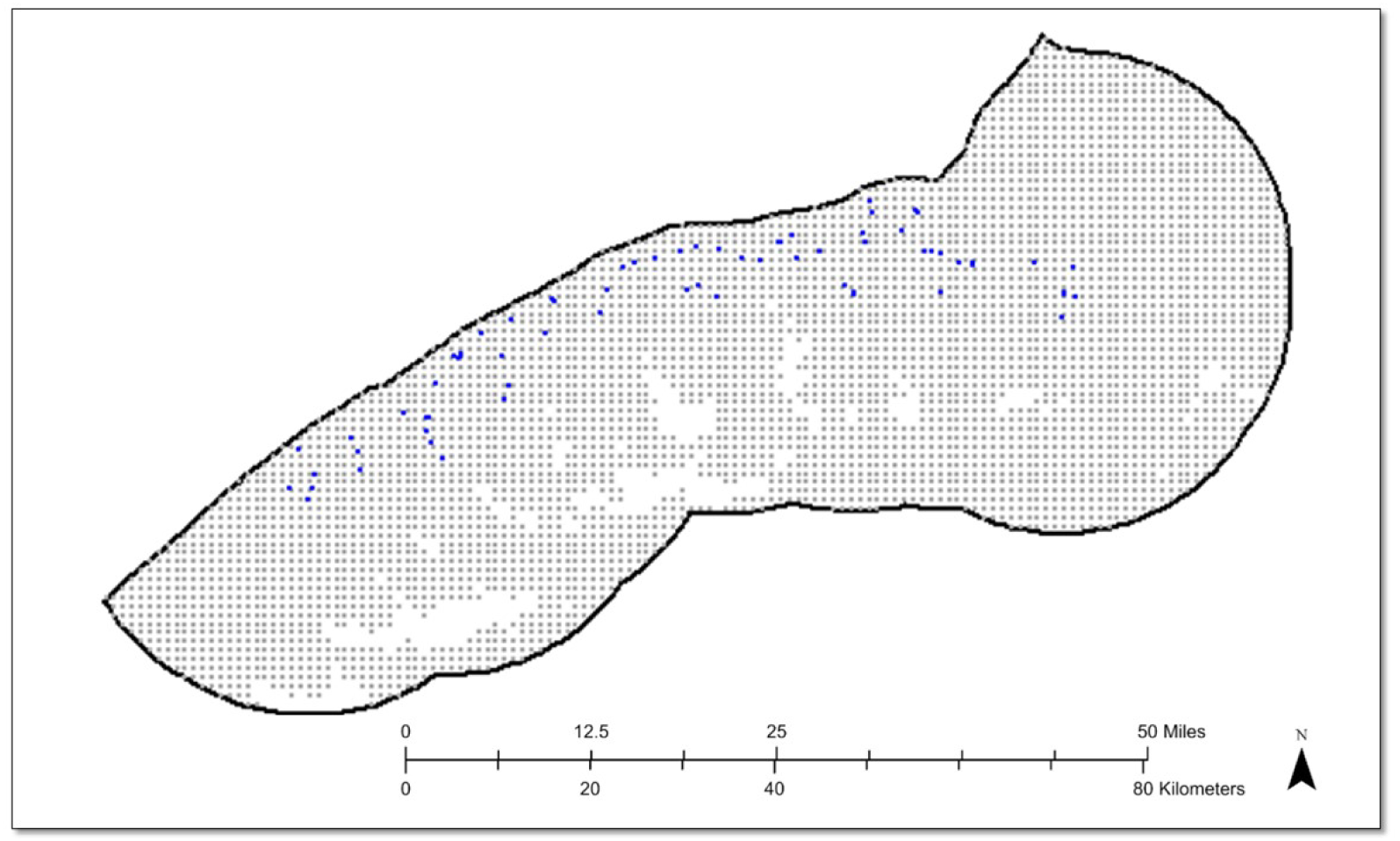
Buffer mask boundary showing camera trap locations (blue), buffer grid (grey), and excluded areas (white). The rigid northern boundary reflects the large river that snow leopards do not cross.

## Results

### Image Classification

Of the 1,054 camera trap encounters recorded during the study periods, 844 were classified as high-quality and included in the final analysis. The remaining 210 encounters were categorized as low-quality and excluded. Among the high-quality encounters, 118 snow leopards were identified, with 42 individuals detected only once and 76 detected more than once (Table 2).

**Table 2.** Summary of high- and low-quality encounters, individual detections, and the number of left, right, and both-sided detections.

| Session | End | Season | High Quality Encounters | Low Quality Encounters | Detected Once | Detected >1 | Left + Both | Right + Both | Left Only | Right Only | Both Sided | Total Individuals |
| --- | --- | --- | --- | --- | --- | --- | --- | --- | --- | --- | --- | --- |
| 1 | 2012 | Winter | 39 | 9 | 7 | 9 | 12 | 11 | 5 | 4 | 7 | 16 |
| 2 | 2013 | Winter | 78 | 24 | 6 | 16 | 14 | 18 | 4 | 8 | 10 | 22 |
| 3 | 2013 | Spring | 46 | 6 | 6 | 7 | 10 | 8 | 5 | 3 | 5 | 13 |
| 4 | 2013 | Spring | 50 | 11 | 8 | 9 | 11 | 13 | 4 | 6 | 7 | 17 |
| 5 | 2013 | Summer | 50 | 16 | 6 | 13 | 16 | 13 | 6 | 3 | 10 | 19 |
| 6 | 2013 | Fall | 36 | 9 | 7 | 8 | 9 | 10 | 4 | 5 | 5 | 14 |
| 7 | 2017 | Fall | 31 | 10 | 12 | 5 | 6 | 14 | 3 | 11 | 3 | 17 |
| 8 | 2018 | Winter | 143 | 24 | 15 | 27 | 27 | 29 | 10 | 12 | 17 | 39 |
| 9 | 2018 | Winter | 119 | 23 | 9 | 21 | 12 | 24 | 2 | 14 | 10 | 26 |
| 10 | 2018 | Spring | 98 | 20 | 13 | 14 | 16 | 19 | 5 | 8 | 11 | 24 |
| 11 | 2019 | Summer | 73 | 53 | 11 | 18 | 17 | 15 | 11 | 9 | 6 | 26 |
| 12 | 2019 | Fall | 81 | 5 | 17 | 14 | 20 | 20 | 10 | 10 | 10 | 30 |
| Totals |  |  | 844 | 210 |  |  |  |  |  |  |  |  |

### Quality Sorting

A total of 61 individual snow leopards were detected from either the left (n = 31) or right (n = 30) flank without corresponding images from the opposite side. These individuals were included in separate left-only and right-only models. Additionally, 52 individuals had both left and right-side captures and were included in both models; however, the corresponding images from the opposite side were excluded to maintain consistency and avoid biases. Five high-quality detections showing only partial views (e.g., tail or top view) were removed from the analysis as they could not be matched to other individuals.

### Spatial Capture-Recapture

Models incorporated left-only + both (with right-sided encounters removed), right-only +both (with left only encounters removed), and full (left, right, both) datasets to evaluate how misidentifications from unresolved side detections influence population size estimates. Single-session models were run independently on each of the 12 sessions and averaged, and multi-session occupancy models were conducted separately for the six sessions in Block 1 (2012-2013) and six sessions in Block 2 (2017-2019). Single-session models averaged across all six sessions per block, and results were compared among data subsets (Tables 3a and 3b).

**Table 3.**
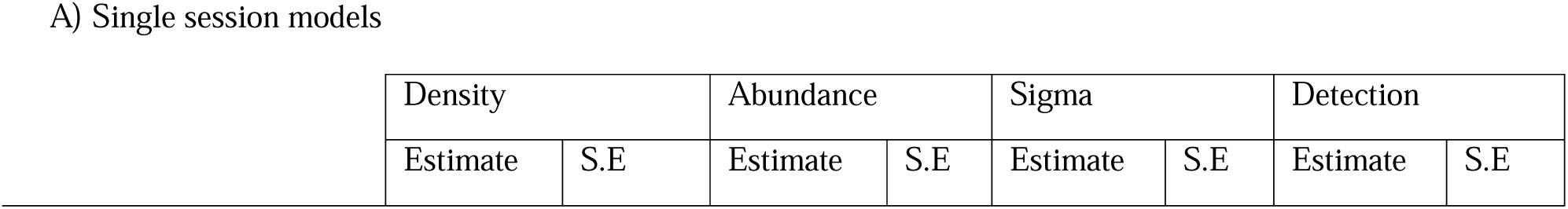
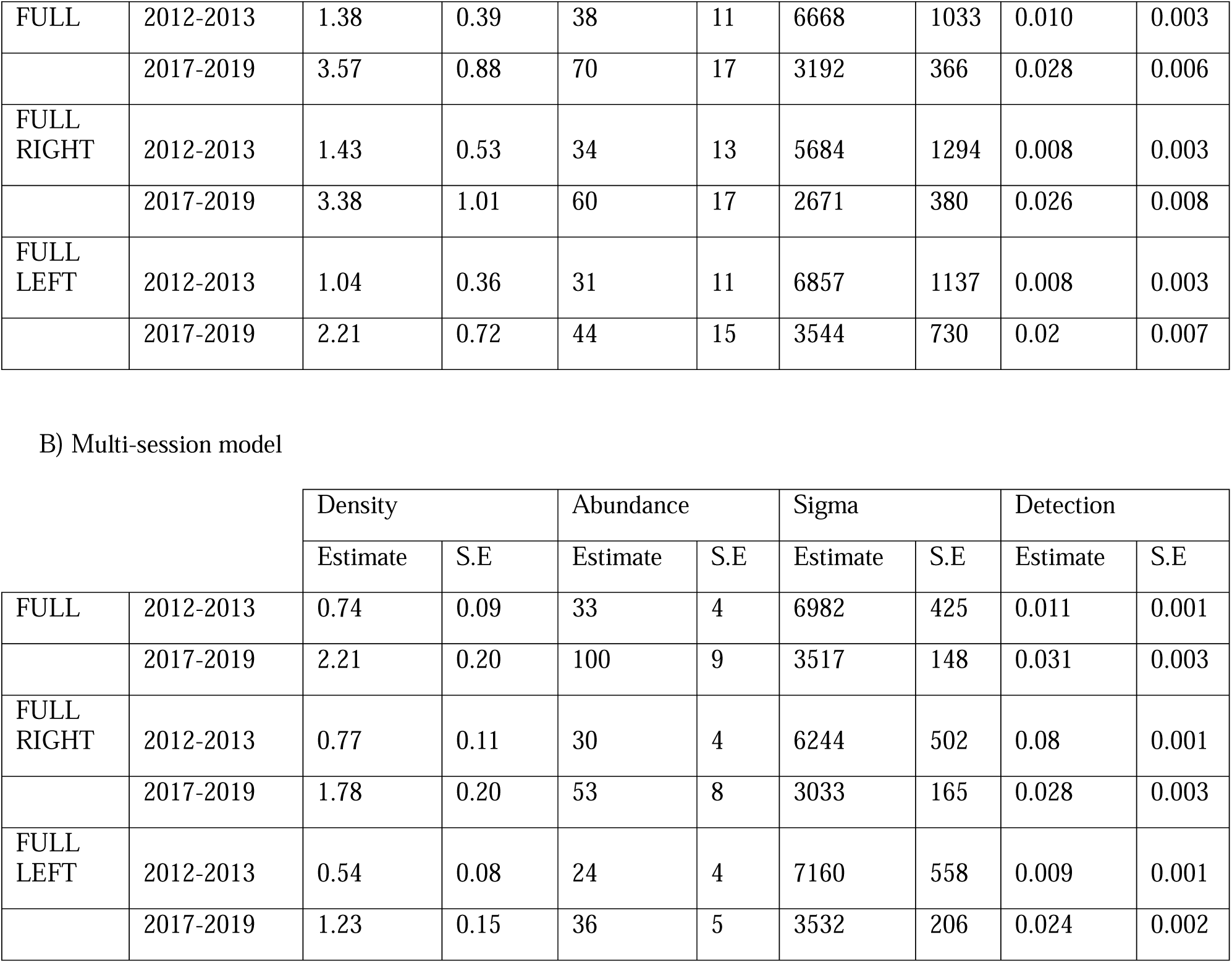
Snow leopard population estimates from (A) single-session averages, and (B) multi-session models, including full, left-only, and right-only datasets. Estimates are provided for density (per 100 km²), abundance, sigma (movement scale parameter in meters), and detection probability.

A) Single session models
| Density |  | Abundance |  | Sigma |  | Detection |  |
| --- | --- | --- | --- | --- | --- | --- | --- |
| Estimate | S.E | Estimate | S.E | Estimate | S.E | Estimate | S.E |

|  |  |  |  |  |  |  |  |  |  |
| --- | --- | --- | --- | --- | --- | --- | --- | --- | --- |
| FULL | 2012-2013 | 1.38 | 0.39 | 38 | 11 | 6668 | 1033 | 0.010 | 0.003 |
|  | 2017-2019 | 3.57 | 0.88 | 70 | 17 | 3192 | 366 | 0.028 | 0.006 |
| FULL<br>RIGHT | 2012-2013 | 1.43 | 0.53 | 34 | 13 | 5684 | 1294 | 0.008 | 0.003 |
|  | 2017-2019 | 3.38 | 1.01 | 60 | 17 | 2671 | 380 | 0.026 | 0.008 |
| FULL<br>LEFT | 2012-2013 | 1.04 | 0.36 | 31 | 11 | 6857 | 1137 | 0.008 | 0.003 |
|  | 2017-2019 | 2.21 | 0.72 | 44 | 15 | 3544 | 730 | 0.02 | 0.007 |

B) Multi-session model
|  |  | Density |  | Abundance |  | Sigma |  | Detection |  |
| --- | --- | --- | --- | --- | --- | --- | --- | --- | --- |
|  |  | Estimate | S.E | Estimate | S.E | Estimate | S.E | Estimate | S.E |
| FULL | 2012-2013 | 0.74 | 0.09 | 33 | 4 | 6982 | 425 | 0.011 | 0.001 |
|  | 2017-2019 | 2.21 | 0.20 | 100 | 9 | 3517 | 148 | 0.031 | 0.003 |
| FULL<br>RIGHT | 2012-2013 | 0.77 | 0.11 | 30 | 4 | 6244 | 502 | 0.08 | 0.001 |
|  | 2017-2019 | 1.78 | 0.20 | 53 | 8 | 3033 | 165 | 0.028 | 0.003 |
| FULL<br>LEFT | 2012-2013 | 0.54 | 0.08 | 24 | 4 | 7160 | 558 | 0.009 | 0.001 |
|  | 2017-2019 | 1.23 | 0.15 | 36 | 5 | 3532 | 206 | 0.024 | 0.002 |

### Single-session model results

In 2012-2013, the single-session model population estimates for snow leopards in Wakhan National Park varied, with the full left model estimating 31 individuals (SE = 11) and a density of 1.04/100 km² (SE = 0.36), while the full right model estimated 34 individuals (SE = 13) and a density of 1.43/100 km² (SE = 0.53). By 2017-2019, population estimates increased significantly: the full left model estimated 44 individuals (SE = 15) with a density of 2.21/100 km² (SE = 0.72), and the full right model estimated 60 individuals (SE = 17) with a density of 3.38/100 km² (SE = 1.01). The full model density estimates rose from 1.38/100 km² (SE = 0.39) in 2012-2013 to 3.57/100 km² (SE = 0.88) in 2017-2019, marking a 159% increase. Comparatively, the density increase for the full right model was 136%, while the full left model showed a 113% increase. Furthermore, the full model’s density estimates were roughly 13% higher in 2012-2013 and 27% higher in 2017-2019 compared to the averages of the left-only and right-only models, illustrating potential overestimation issues linked to unresolved bilateral asymmetry. The abundance estimates using the full model also showed a notable rise, from 38 individuals (SE = 11) in 2012-2013 to 70 individuals (SE = 17) in 2017-2019, an 84% increase. The full right model’s abundance estimates increased from 34 (SE = 13) to 60 (SE = 17), and the full left model rose from 31 (SE = 11) to 44 (SE = 15), representing increases of 76% and 42%, respectively. When compared to the right and left models’ averages, the full model abundance estimates were approximately 16% higher in 2012-2013 and 29% higher in 2017-2019, emphasizing the need for advanced modeling approaches to minimize overestimation biases.

### Multi-session models results

In multi-session models, population estimates increased significantly from 24 individuals (left-only) and 30 individuals (right-only) in 2012-2013, with densities of 0.54/100 km² (SE = 0.08) and 0.77/100 km² (SE = 0.11), respectively, to 36 individuals (left-only) and 53 individuals (right-only) in 2017-2019, with densities of 1.23/100 km² (SE = 0.15) and 1.78/100 km² (SE = 0.20). The full model density estimate rose from 0.74/100 km² (SE = 0.09) in 2012-2013 to 2.21/100 km² (SE = 0.20) in 2017-2019, representing a 199% increase. The full right model’s density increased by 131%, from 0.77/100 km² (SE = 0.11) to 1.78/100 km² (SE = 0.20), while the full left model saw a 128% rise, from 0.54/100 km² (SE = 0.08) to 1.23/100 km² (SE = 0.15). The full model density estimates were approximately 16% higher in 2012-2013 and 32% higher in 2017-2019 compared to the average of the right and left models, suggesting potential biases linked to model assumptions. Abundance estimates from the full model increased from 33 individuals (SE = 4) in 2012-2013 to 100 individuals (SE = 9) in 2017-2019, an increase of 203%. These full model abundance estimates were about 13% higher in 2012-2013 and 39% higher in 2017-2019 compared to the averages of the right-only and left-only models, emphasizing the necessity for refined modeling techniques to ensure accurate population assessments.

## Discussion

Our findings suggest that the snow leopard population in the Wakhan Corridor increased between 2012-2013 and 2017-2019. To our knowledge, this is the first study using camera trap data from two distinct periods to assess snow leopard population size and density changes over time. Previous population estimates for the entire Wakhan National Park (district) were based on extrapolations (Moheb and Paley 2016), but our study, focusing on a smaller research area, yielded more refined results. Between 2012-2013 and 2017-2019, single-session and multi-session models showed significant increases in snow leopard density and abundance. Multi-session models similarly indicated a density increase.

While snow leopards live up to 20 years in captivity (Hunter and Barrett 2020), their lifespan in the wild is likely 10–13 years (Nigam et al. 2012). Snow leopards breed seasonally, with litters of one to four cubs born in spring.

Cubs are weaned after two months, but juveniles stay with the mother for two years and can start breeding at 33 months (Mallon et al. 2016; McCarthy et al. 2017; Hunter and Barrett 2020). Population growth such as that identified by this study in increases in multiple captured individuals, ∼0-12 more individuals across single and multi-session models, would be possible in 5-7 years. If the sex ratio is 50%, ∼15 females bred 2-3 cubs between 2012-2013, we could observe an increase in snow leopards by 2017-2019. Birth rates contribute to a growing population, but immigration from other areas, such as Afghanistan and Tajikistan (Haqiq thesis), has also likely played a role (Johansson et al. 2018). Despite some limitations imposed by the five-year gap between two years of data, this study’s findings indicate a population size increase.

Detection probabilities improved across all models, suggesting better survey effectiveness over time. Multi-session models provided more reliable yearly estimates than single-session data, which may not fully capture year-round population dynamics. We explored the use of multi-session models with consecutive sessions, thus incorporating individuals across sessions, similar to other studies (Wang et al. 2018; Gabriele-Rivet et al. 2020). Although this method technically violated temporal independence, the literature suggests overlapping individuals across sessions can increase precision, although with less exact standard errors (Sutherland et al. 2019; Efford 2021a). Multi-session models provide more reliable yearly estimates than averaging single-session data. Future research could explore open population spatial capture-recapture models (Augustine et al. 2019), SPIM (Augustine et al., 2018), or pursue the development of an open population SPIM model, acknowledging that such a model has yet to be established.

To assess variations and potential overestimation, we analyzed population estimates using three data parameterizations—full, left-only, and right-only datasets. Our results revealed a significant impact of bilateral asymmetry on estimates. For single-session models, the full model density estimates were higher compared to the average of the left-only and right-only models. Similarly, the full model density estimates were also much higher for multi-session models. These findings re-emphasize the need to consider unresolved bilateral detections in camera trap data to avoid inflated population estimates. Previous snow leopard studies that did not account for such asymmetry may have overestimated snow leopard populations. While our study does not assert exact densities, it highlights a high potential for misidentification. Therefore, estimates reported in the literature that did not account for left and right flanks may not accurately reflect the true data.

AI-assisted individual identification and expert validation played a critical role in reducing misclassification bias in this study. This study is the first to estimate snow leopard density using AI-assisted individual identification and an expert observer qualifying the AI matches. Given the demonstrated ability of snow leopard algorithms to perform well on high-quality datasets (Bohnett et al. 2023a), Whiskerbook’s accuracy in identifying individual snow leopards was important for managing the large dataset. AI algorithms combined with user expertise significantly increase snow leopard detection accuracy and efficiency (Bohnett et al. 2023b).

We discovered that single-capture individuals constituted a significant portion of our dataset (Table 2). These individuals nearly doubled population estimates in 2017-2019, suggesting that they use the study area less frequently or may be transient. Since AI has vastly improved our capabilities for achieving accuracy in camera trap studies, the issues with singleton encounters being misclassifications were found to have been substantially improved (Bohnett et al. 2023a). However, recent studies have addressed these single “ghost” encounters by a different parameterization (Kodi et al. 2024).

Long-term datasets spanning at least three years could help distinguish transient or dispersing snow leopards from resident individuals despite challenges related to population closure. Such datasets may identify individuals lacking permanent home ranges, including young, aging, or seasonally shifting snow leopards, and those with large adjacent home ranges. GPS collaring studies (Johansson et al. 2018; Rosenbaum et al. 2023) show that snow leopards range significantly in distances and seasonally adjust home ranges, emphasizing the importance of analyzing site use. Their solitary, territorial nature includes sex-specific seasonal trends, with females expanding ranges for reproduction and food and males adjusting for mating or prey movement (Johansson et al. 2018; Rosenbaum et al. 2023).

Long-term population trends regarding environmental changes, conservation efforts, and emerging threats should also be evaluated. Shifting conservation priorities, climate change, and human-wildlife conflict influence snow leopard populations (Hoffmann et al. 2010, 2011; Li et al. 2016; Shen 2020; Hoffmann and Beierkuhnlein 2020). Except for a small number of studies (Taubmann et al. 2016; Ghoshal et al. 2019), there is little evidence of snow leopard population changes across adequate periods to inform population estimates, assess conservation activities or threats, and determine their consequences (McCarthy et al., 2017, Taubmann et al., 2016). From 1990 to 2010, sign surveys and expert interviews across 14,000 km^2^ of Kyrgyz Alay suggested a 39% decrease in this local snow leopard population (Taubmann et al. 2016). Another Himachal Pradesh study evaluated snow leopard site use using recall-based expert interviews and reported no population change from 1985-1992 and 2008-2012 (Ghoshal et al. 2019). Another study estimated recruitment and survival using 2009–2012 camera trapping data and found a stable snow leopard population in Mongolia’s Tost mountains (Sharma et al. 2014). While the snow leopard population in Wakhan appeared to increase between 2012 and 2017, it remains unclear if this trend has continued following the collapse of the elected government in 2021 and the disruption of wildlife protection structures. Given recent signs of a declining prey base due to uncontrolled poaching, conducting a new study as soon as conditions permit is crucial to assess the current trends impacting the snow leopard population. Understanding the current trends impacting the snow leopard population is critical for informing conservation strategies and addressing emerging threats.

### Conservation implications

The increasing snow leopard population in the Wakhan Corridor suggests that conservation efforts carried out between 2012 and 2017 yielded positive results. The establishment of Wakhan National Park in 2014, threat mitigation, and community-based conservation efforts contributed to these positive results (Moheb et al., 2022, UNDP, 2021, WCS Afghanistan, 2021). In 2021, Afghanistan experienced significant political changes that have inevitably affected the conservation efforts undertaken in the country by the previous government. The new Islamic Emirate of Afghanistan appears motivated to conserve the country’s unique flora and fauna (NBSAP revised 2023), and it is hoped that this goodwill will translate into effective conservation actions.

## Conclusions

Effective wildlife monitoring and conservation rely on accurate assessments of species populations to track trends over time. This study applied SCR-based camera trapping methodologies, integrating AI for individual identification and modeling frameworks to address bilateral asymmetry in snow leopards. By employing single-and multi-session open spatial capture-recapture (oSCR) models, we aimed to provide the most accurate population estimates. These methods can be adapted to other carnivore species facing similar challenges. This study offers a comprehensive analysis of snow leopard population dynamics in the Wakhan Corridor 2012-2013 and 2017-2019 by employing camera trapping methodologies combined with spatial capture-recapture (SCR) models and AI-assisted individual identification. Our findings provide new insights into snow leopard population trends over time, addressing previous gaps in knowledge. Our use of AI-assisted identification through Whiskerbook, paired with expert validation, is believed to have improved the accuracy of individual identification for the species (Johansson et al. 2020; Bohnett et al. 2023b). This methodological advancement is crucial for large-scale wildlife monitoring efforts. This study found a significant increase in density and abundance between 2012-2013 and 2017-2019. We achieved more accurate estimates by applying advanced spatial capture-recapture models and AI-assisted individual identification, highlighting an increase in density and abundance on average between the right-only and left-only models over the study period. These results underscore the effectiveness of integrating technological advancements in wildlife monitoring, which can offer improved insights into population dynamics. Our findings also emphasize the importance of addressing bilateral asymmetry in camera trap data, as unresolved side detections can lead to inflated population estimates. By incorporating both left-only and right-only datasets, we identified a substantial overestimation potential, with the full model estimates being significantly higher. This insight is critical for future studies and conservation efforts, as it clarifies population size and trends. Overall, the observed population growth suggests that conservation efforts and environmental conditions in the Wakhan Corridor may support the recovery of snow leopards. However, continued monitoring and long-term studies are essential to account for potential changes in population dynamics due to climate change, human-wildlife conflict, and habitat alterations. The study’s approach and findings contribute valuable knowledge to the field of wildlife conservation and provide a foundation for future research and management strategies.

## Financial interests

The authors have no relevant financial or non-financial interests to disclose.

Conflicts of interest None

Ethical standards European Union project “Improving participatory management and efficiency of rangeland and watershed focusing on Wakhan, Yakawlang, Kahmard and Saighan Districts (Contract ACA/2018/399-742)” ethical standards for animal data collection were met. UNDP GEF grants 00076820/0088001/5038 and 00105859/00106885/5844. Segré Foundation grant “Transboundary Mountain Monarch Conservation in Afghanistan and Pakistan".

Data availability Data are available upon request from the Whiskerbook.org web portal. All data are available online.

## Acknowledgments

We appreciate Afghanistan, USAID, GEF/UNDP, Fondation Segré, and the EU for allowing the Wildlife Conservation Society to perform the study programs. The author’s opinions are their own and do not necessarily reflect EU policy. Thanks to Dr. Richard Paley, Country Director of WCS Afghanistan, for his inspiring leadership, Ali Madad Rajabi for supervising camera trapping, Shanbe Shirzad, Ayan Big, Aziz Big, Karmal for camera trap deployment, and the WCS Kabul and Wakhan teams for their invaluable support. We also thank the Afghan people for their hospitality and fieldwork support. Finally, we thank the anonymous reviewers for their time and effort checking the work.

